# One of these things is not like the others: Theta, beta, & ERP dynamics of mismatch detection

**DOI:** 10.1101/2025.07.11.664390

**Authors:** Josset B. Yarbrough, Lingxiao Shi, Kaustav Chattopadhyay, Robert T. Knight, Elizabeth L. Johnson

**Author notes:** Corresponding authors. E-mail addresses (J.B. Yarbrough), (E.L. Johnson).

## Abstract

Working memory (WM) enables the detection of mistakes by permitting one to notice when sensory input is *mismatched* to their internal prediction. Prior studies support the role of frontal midline theta activity, with an overlapping N200 event-related potential (ERP), as a mechanism for comparing incoming sensory stimuli to the internal model. Additionally, posterior low-beta activity has been proposed as a mechanism for processing incoming sensory stimuli in WM. However, it is unknown whether frontal midline theta activity and the N200 support mismatch detection, or whether posterior low-beta activity extends from sensory processing to detecting a mismatch between sensory input and the internal model. Here, we reveal that frontal midline theta supports mismatch detection and explains individual WM performance. Unexpectedly, instead of the N200, results show a positive slow wave ERP overlapping with the frontal midline theta mismatch response. Results additionally indicate a late posterior low-beta response persisting from stimulus presentation into the post-stimulus delay. Our findings establish frontal midline theta as a marker of successful mismatch detection, challenge the domain-general role of the N200 in error detection, and support theories linking posterior low-beta to processing incoming sensory stimuli.

## Introduction

Working memory (WM) governs the rapid storage, maintenance, manipulation, and retrieval of incoming information. Effective WM can detect stimulus mismatches – wherein external stimuli deviate from one’s internal prediction, like noticing a band play the wrong note while performing a familiar song – and adjust performance. Mismatch detection is similar to error detection because, instead of responses being generated from processing unexpected internal actions, mismatch responses are generated from processing unexpected external sensory stimuli. Prior studies of WM and control present converging evidence that novelty, error, and conflict elicit similar frontal midline theta activity (FMθ; 4-8 Hz)^1^. However, it is unclear whether FMθ supports mismatch detection during WM as information is processed in real time, and whether other electrophysiological responses to mismatches may be related to or distinct from the FMθ response. Event-related potentials (ERPs) including the N200 and the P300 have also been linked to novelty, error detection, and conflict monitoring^1,2^. Additionally, maintaining sensory information in WM has been linked to posterior oscillations in the low-beta range (Pβ1; 12-20 Hz)^3,4,5^. There remains a gap in the literature on the electrophysiological mechanisms of mismatch detection during WM.

A ubiquitous spectral signature of cognitive control across rodents and primates is FMθ^1^. Substantial evidence supports the role of FMθ in top-down control during events that challenge the brain’s internal prediction model such as novelty, error, and conflict. For instance, increased FMθ power has been observed both in response to an “oddball” stimulus and to externally presented feedback following an incorrect behavioral output^6^. The classic “oddball task,” where participants listen to a series of the same tones mixed intermittently with oddball (i.e., different) tones, demands WM to detect a mismatch between the external sensory stimulus and one’s internal prediction model. Human scalp electroencephalography (EEG) studies have consistently reported N200 and P300 ERPs, specifically the P3a frontal response to external stimuli, in response to the oddball stimulus^2,7^. These oddball ERPs share frontal topography but differ in purpose; the P3a is associated with attentional capture by novel or unexpected stimuli, whereas the N200 is associated with error detection and stimulus maintenance^1,2^. In addition, the parietal P3b is associated with updating the brain’s internal model following an oddball stimulus^2^. The N200 and FMθ are both thought to be generated by the anterior cingulate cortex (ACC), and the N200 peak during the processing of novelty and conflict shares similar timing and topography as the FMθ response^6^. FMθ has also been observed during the manipulation of letter sequences in WM, demonstrating that it is involved in top-down control and updating the contents of WM^8^. Increased FMθ has also been correlated with increased WM load and decreased retention of unfamiliar shapes^9^. Accordingly, FMθ has been proposed as an electrophysiological substrate of the “central executive” (the attention controller) in the classic three-component model of WM^1,6^ which also includes the “phonological loop” (a limited store of speech related information) and the “visuospatial sketchpad” a limited store of visual and spatial information)^1,6,10^. One might predict FMθ to support mismatch processing during WM and predict behavioral outcomes.

In addition to control, essential to WM is the encoding and maintenance of sensory information. To effectively process sensory inputs, there needs to be a temporary space that allows for control mechanisms to process the incoming stream of information. Baddeley coined this space the “episodic buffer,” and included it as the fourth component in an update to the classic three-component model of WM^11^. The episodic buffer is thought to be a higher-level space that has the capacity to bind multiple features of sensory information and is accessible to the central executive^12^. Pβ1 is proposed to be a substrate of the episodic buffer by enabling long-range synchrony to combine sensory information with frontal control decisions to update one’s internal predictive model^3^. Increased Pβ1 has been observed during the delay periods of visuospatial and Sternberg WM tasks, supporting Pβ1 as a mechanism by which sensory information is processed to achieve behaviorally relevant goals^5^. Pβ1 has also been observed during the processing of auditory oddball stimuli, suggesting that it is also involved in detecting novel or unexpected stimuli as mismatches^13^. By processing features of sensory stimuli, Pβ1 provides another candidate electrophysiological mechanism for the detection of mismatches during WM through comparison of external sensory information to one’s internal model.

We designed a sequential WM delayed match-to-sample (WM-DMS) task to establish electrophysiological mechanisms of mismatch detection during WM (**Fig. 1**). On each trial, participants view a sample sequence of three different shapes in specific spatiotemporal positions, followed by a post-sample delay and one of four test sequences. The test sequence is either an exact match, or two of three stimuli are mismatched on the identity, temporal, or spatial dimension. After the post-test delay, participants indicate whether the test sequence is a match or a mismatch to the sample sequence. We collected scalp EEG data to define ERP and spectral signatures of mismatch detection as each stimulus is presented in the test sequence.

**Figure 1.**
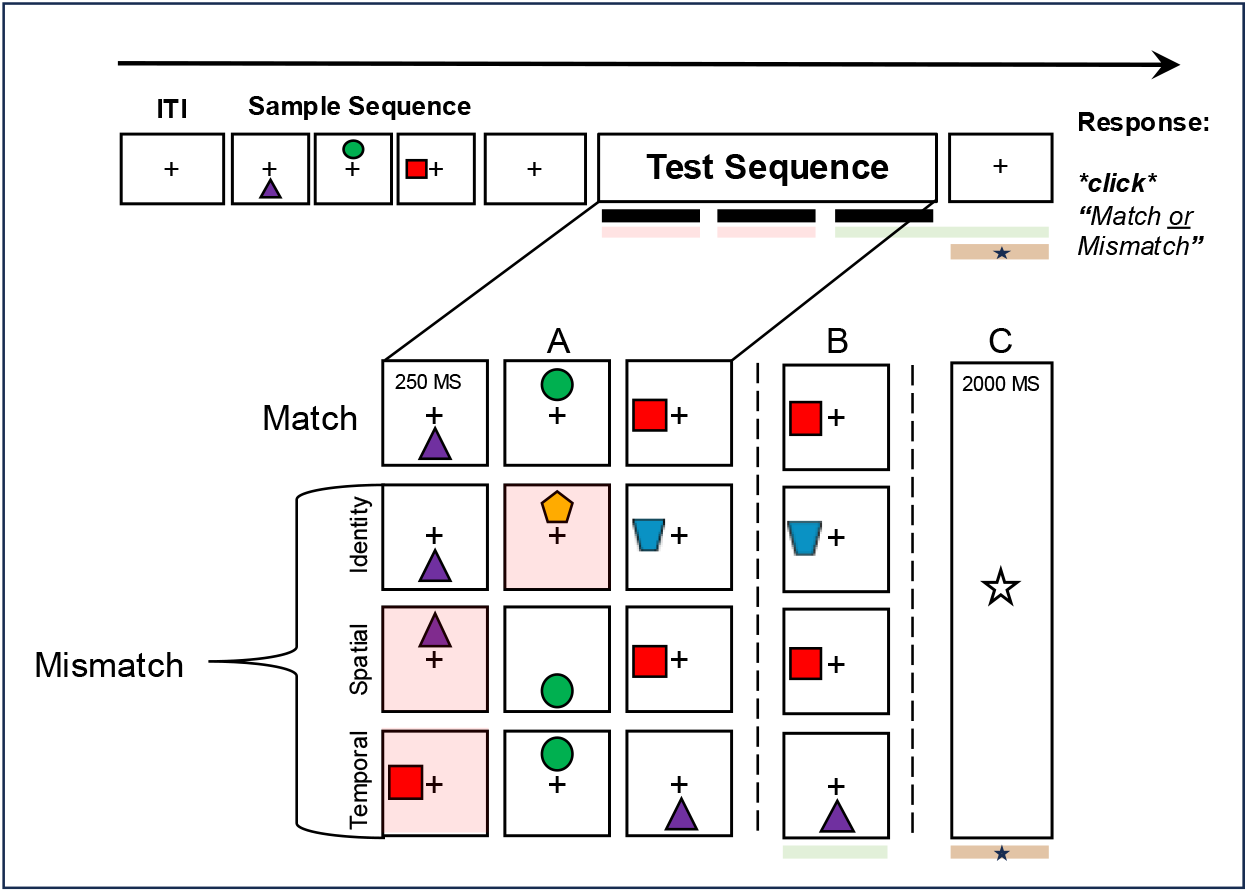
WM-DMS paradigm. On each trial, participants viewed a sample sequence of three shapes in specific spatiotemporal positions (0.25 s stimulus, 0.25 s interstimulus interval/ISI), followed by a 2-s delay and a test sequence with identical timing to the sample sequence. The test sequence matched the sample sequence exactly or two out of three stimuli were mismatched on one dimension (identity, spatial, or temporal). ITI, intertrial interval. **A)** Trials realigned to the first mismatched stimulus, which occurred first or second in the test sequence (pink), fully counterbalanced across conditions. Matched test sequences were submitted to the same realignment procedure to account for any position effects. There were three mismatch types: identity (i.e., 2 new shapes), spatial (i.e., location of 2 shapes swapped), and temporal (i.e., order of 2 shapes swapped). **B)** No-star trials aligned to the ISI between the second and third test stimuli, extending through the post-test delay period (green) to examine persistent activity. **C)** Star trials realigned to the star onset during the post-test delay (brown) to examine latent activity. The epoch was extended -1 s and +0.85 s from the star onset.

To identify these signatures, we aligned the EEG data from correct trials to the onset of the first mismatched stimulus in each test sequence and compared mismatched to matched test sequences (see **Fig. 1A**). We hypothesized that the mismatch would elicit N200 and P300 ERPs as well as increased FMθ and Pβ1 activity. Furthermore, we hypothesized that FMθ would predict task performance measures, that is, behavioral accuracy rate and response time (RT).

After testing hypotheses, we explored whether mismatch responses might continue to be represented in a persistently active or latent state by selectively “pinging” the brain during the delay^14–24^.On half of the trials, we briefly flashed a star in the center of the screen at a randomly jittered time during the post-sample and post-test delays, and participants were told to ignore it. Flashing the star, a task-irrelevant stimulus, reactivates stimulus-specific activity patterns during short-term synaptic plasticity and can thus elicit measurable responses related to otherwise latent WM representations^18^ .To identify potential latent representations, we realigned the EEG data from these trials to the onset of the star during the post-test delay and again compared mismatched to matched test sequences (see **Fig. 1C**). To identify potential persistent representations, we analyzed the post-test delay phase on the other half of trials without any realignment (see **Fig. 1B**). We considered mismatch responses during the no-star and star delay periods as indicative of persistent and latent representations, respectively. Evidence of persistent activity would be indicated by the temporal extension of a mismatch response from the test sequence into the post-test delay^25^, supporting the maintenance of match or mismatch information through the post-test delay and up to the RT. By contrast, evidence of latent activity would be indicated by a mismatch response following the star^18^.

## Results

### Behavior

Participants were selected on the basis of above-chance behavioral performance across mismatch and match trials and, accordingly, our sample performed well (M ± SD, 0.85 ± 0.12; one-sample t(34) = 40.35, p = 3×10^−30^). RT was on average 1.78 ± 0.52 s. A significant negative correlation between accuracy rate and RT (rho = -0.34, p = 0.046) demonstrated that individuals with superior performance tended to respond faster. There were no significant differences in accuracy rate or RT between star and no-star trials (p > 0.34).

### Mismatches elicit positive slow wave ERPs in frontal midline channels

To identify ERP signatures of mismatch detection, we compared mismatched and matched test sequences locked to the onset of the first mismatched stimulus. A significant cluster-corrected t-test revealed mismatch > match ERP effects in frontal midline channels (p = 0.0001, Cohen’s d = 1.57; **Fig. 2A**). Effects were characterized by increased voltage on mismatch trials that was sustained from 0.35-0.85 s, starting during the ISI following the first mismatched stimulus and ending during the ISI following the next stimulus. This type of sustained positive ERP has been termed the positive slow wave (PSW). The PSW effect was not significantly correlated with behavioral accuracy rate or RT. No other ERP effects reached significance.

**Figure 2.**
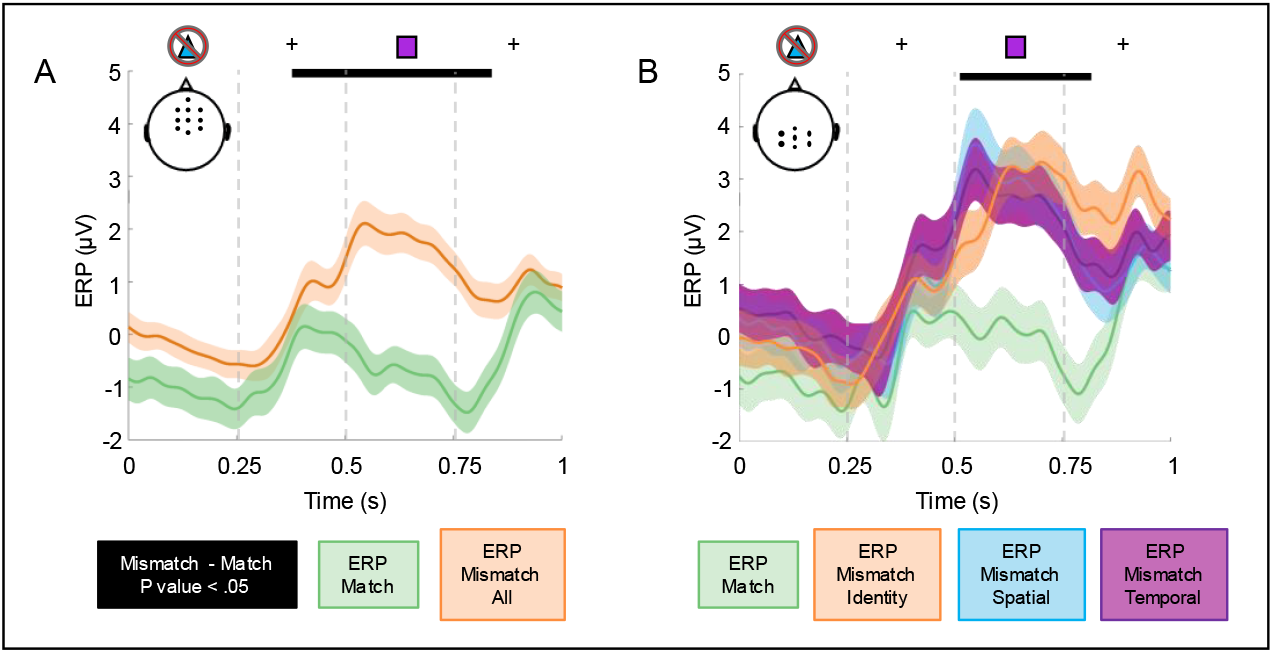
Frontal midline PSW ERP mismatch effect. **A)** Grand mean ERPs across channels with significant mismatch (orange) > match (green) effects (inset), determined by a t-test. Shading represents SEM. Time = 0 indicates the onset of the first mismatched stimulus. The crossed-out triangle represents the first mismatched stimulus in the test sequence and the square represents the next stimulus, separated by fixation ISIs. Grey dotted lines represent stimulus onsets and offsets. The black bar indicates time points of significant effects. **B)** Same as (A) for an F-test examining all four conditions.

In addition, we compared all four types of test sequence without collapsing across the three mismatch conditions. A significant cluster-corrected F-test confirmed mismatch > match ERP effects (p = 0.023), with no significant differences between mismatch conditions (**Fig. 2B**).

### Mismatches elicit increased theta activity and phase clustering in frontal midline channels

We applied the same approach to identify spectral power and ITPC signatures of mismatch detection. A significant cluster-corrected t-test revealed mismatch > match effects in FMθ power (p = 0.0001, Cohen’s d = 1.29; **Fig. 3A**). Effects were characterized by increased power from 2.5-15.0 Hz on mismatch trials that was focused in the θ range and sustained from 0.25-0.75 s, starting at the offset of the first mismatched stimulus and ending at the offset of the next stimulus. The FMθ power effect correlated positively with subsequent behavioral accuracy (rho = 0.53, p = 0.0027; **Fig. 3B**).

**Figure 3.**
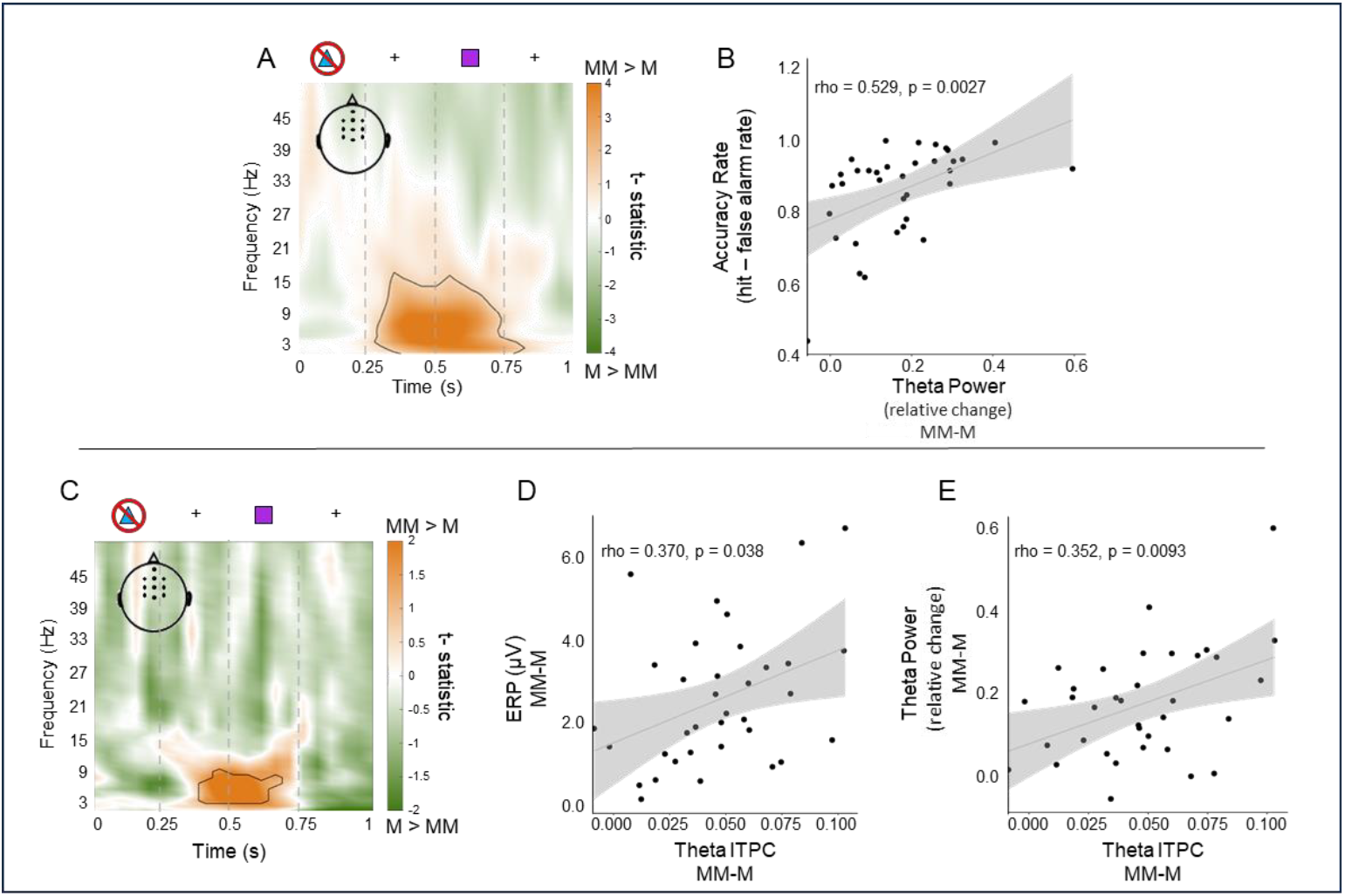
FMθ power and ITPC mismatch effects. **A)** Time-frequency representation of t-statistics on power across frontal channels with significant mismatch > match effects (inset). Time = 0 indicates the onset of the first mismatched stimulus. The outline indicates time-frequency points of significant effects, focused in the θ range. The crossed-out triangle represents the first mismatched stimulus in the test sequence and the square represents the next stimulus, separated by fixation ISIs. Grey dotted lines represent stimulus onsets and offsets. MM, mismatch; M, match. **B)** Significant positive correlation between FMθ power mismatch effect in (A) and behavioral accuracy rate. Dots represent individual participant mismatch – match differences. Shading represents SEM. **C)** Same as (A) for ITPC. **D)** Significant positive correlation between FMθ ITPC mismatch effect in (C) and the frontal midline PSW ERP effect (see **Fig. 2**), same conventions as (B). **E)** Significant positive correlation between FMθ TPC mismatch effect in (C) and FMθ power mismatch effect in (), same conventions as (B).

A significant cluster-corrected t-test likewise revealed mismatch > match effects in FMθ TPC (p = 0.0001, Cohen’s d = 1.6 ; **Fig. 3C**). Effects were characterized by increased ITPC from 2.5-9.0 Hz that was focused in the θ range and sustained from 0.40-0.70 s, starting during the ISI following the first mismatched stimulus and ending during the presentation of the next stimulus. The FMθ TPC effect correlated positively with both the frontal midline PSW ERP effect (rho = 0.37, p = 0.038; **Fig. 3D**) and the FMθ power effect (rho = 0.35, p = 0.0093; **Fig. 3E**).

### Mismatches elicit low-beta activity and reduce phase clustering in posterior midline channels

The same cluster-corrected t-tests of spectral power and ITPC also revealed a distinct set of posterior midline effects. There was a significant mismatch > match effect in Pβ1 power (p = 0.005, Cohen’s d = 1.11; **Fig. 4A**). Effects were characterized by increased power from 5.0-30.0 Hz on mismatch trials that was focused in the β1 range and sustained from 0.75-1.0 s, starting at the offset of the stimulus following the first mismatched stimulus and extending into the ISI/delay. The Pβ1 power effect was not significantly correlated with behavior.

**Figure 4.**
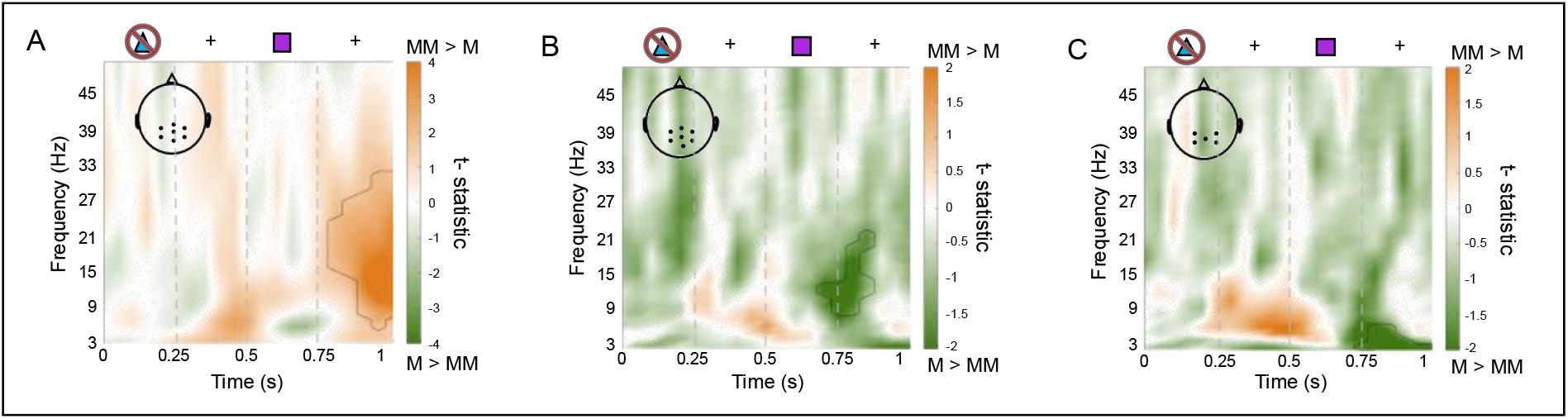
Posterior midline power and ITPC mismatch effects. **A)** Time-frequency representation of t-statistics on power across posterior channels with significant mismatch > match effects (inset). Time = 0 indicates the onset of the first mismatched stimulus. The outline indicates time-frequency points of significant effects, focused in the β1 range. The crossed-out triangle represents the first mismatched stimulus in the test sequence and the square represents the next stimulus, separated by fixation ISIs. Grey dotted lines represent stimulus onsets and offsets. MM, mismatch; M, match. **B)** Time-frequency representation of t-statistics on ITPC across posterior channels with significant β1 mismatch match effects (inset), same conventions as (A). **C)** Same as (B) for delta effects.

There were significant mismatch < match effects in both Pβ1 (p = 0.0001, Cohen’s d = 1.50; **Fig. 4B**) and posterior midline delta (p = 0.014, Cohen’s d = 1.44; **Fig. 4C**) ITPC. The Pβ1 effects were characterized by decreased ITPC from 7.0-21.0 Hz on mismatch trials that was focused in the β1 range and sustained from 0.70-0.85 s, starting during the presentation of the stimulus following the first mismatched stimulus and ending soon after. The delta effects were characterized by decreased ITPC from 2.5-6.0 Hz on mismatch trials that was sustained from 0.75-1.0 s, starting at the offset of the stimulus following the first mismatched stimulus and extending into the ISI/delay. The posterior midline ITPC effects were not significantly correlated with behavior or other posterior effects.

### Mismatch effects persist into the delay in posterior midline channels

Having revealed ERP and spectral signatures of mismatch detection, we explored whether mismatch responses might continue to be represented in a persistently active or latent state. To identify potential persistent representations, we compared mismatch and match ERP, power, and ITPC data on no-star trials from the offset of the second stimulus in the test sequence through the post-test delay without realignment (see **Fig. 1B**). We operationalized persistent representation as an effect that persisted from presentation of the test sequence into the post-test delay^25^. Because posterior midline effects in spectral power and ITPC were identified well after the offset of the first mismatched stimulus (see **Fig. 4**), we anticipated that they would persist into the post-test delay. A significant cluster-corrected t-test revealed mismatch > match effects in Pβ1 power (p = 0.0001, Cohen’s d = 1.09; **Fig. 5A**). Effects were characterized by increased power from 8.0-30.0 Hz on mismatch trials that was focused in the β1 range and sustained from -0.5-0.75 s from delay onset, starting at the offset of the second stimulus and persisting into the delay. In addition, a significant cluster-corrected t-test revealed mismatch < match effects in Pβ1 and delta ITPC (p = 0.0001, Cohen’s d = 2.39; **Fig. 5B**). Effects were characterized by decreased ITPC from 2.5-22.0 Hz on mismatch trials that was sustained from - 0.5-0.5 s, starting at the offset of the second stimulus and persisting into the delay. The β1 effect was time-locked to the offset of each stimulus and clustered with a sustained delta effect. There were no significant ERP effects.

**Figure 5.**
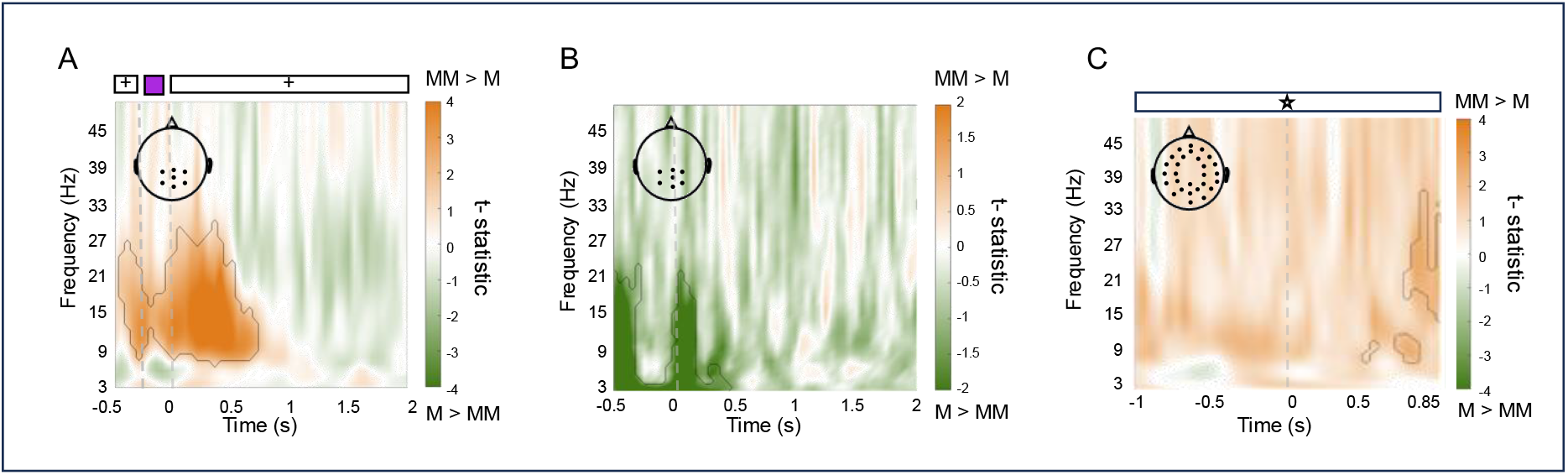
Persistent and latent mismatch effects. **A)** Time-frequency representation of t-statistics on power across channels with significant mismatch > match effects on no-star trials (inset). Time = 0 indicates the onset of the post-test delay. The outline indicates time-frequency points of significant effects, focused in the β1 range and persisting into the delay. The square represents the last stimulus in the test sequence, between the second fixation ISI and delay. Grey dotted lines represent stimulus onset and offset. MM, mismatch; M, match. **B)** Time-frequency representation of t-statistics on ITPC across channels with significant mismatch < match effects on no-star trials (inset), same conventions as (A). **C)** Time-frequency representation of t-statistics on power across channels with significant mismatch > match effects on star trials (inset). Time = 0 indicates the onset of the star during the post-test delay. The outline indicates time-frequency points of significant effects, largely focused in the beta range and time-locked to the star presentation. MM, mismatch; M, match.

### Pinging the brain reveals latent mismatch effects

To identify potential latent representations of mismatch responses, we compared mismatch and match ERP, power, and ITPC data from the post-test delay on star trials locked to the onset of the star (see **Fig. 1C**). We operationalized latent representation as an effect that was time-locked to the star presentation^18,25^. A significant cluster corrected t-test revealed mismatch > match effects in whole-scalp alpha-beta power (p = 0.006, Cohen’s d = 1.43; **Fig. 5C**). Effects were characterized by increased power from 5.0-35.0 Hz on mismatch trials that was largely focused in the beta range between 0.50 and 0.85 s from star onset, starting after the offset of the star and extending through the delay. There were no significant ERP or ITPC effects.

## Discussion

Here we reveal dissociable frontal and posterior electrophysiological signatures of mismatch detection. In line with our hypothesis and prior literature, we found that mismatches elicit FMθ activity during real-time mismatch detection in WM, and that the FMθ power response to mismatches predicts superior WM accuracy. Our findings support dominant models stating that increased FMθ indicates the need for more executive control^1^ and establish the role of FMθ in mismatch detection. Our findings further support FMθ as an electrophysiological substrate of the central executive in the classic three-component model of WM^1,6,10^.

Together with increased FMθ activity, we hypothesized that we would see an N200 ERP mismatch response in the same frontal midline topography, as well as a posterior P300 response. Our results did not support this hypothesis. Instead of an N200 response or other negative components such as the error-related or mismatch negativity^1^, mismatched stimuli elicited a frontal midline PSW response that correlated with FMθ ITPC. We did not observe a significant posterior P300 response, potentially due to our task having more mismatched than matched trials, making the mismatch novel but not unexpected *per se*, thus decreasing prediction error effects. However, we note that a more anterior P300 response may have been part of the frontal midline PSW response, consistent with literature on the oddball task^2^. The PSW ERP is characterized by a late, sustained positive potential in a central topography, and has been related to arousal and processing novel stimuli^26^. The PSW ERP has also been observed following the detection of an auditory target in a cognitively demanding task ^27^. Our ERP findings suggest that mismatch detection elicits a domain-general novelty response associated with increased cognitive demand.

We next hypothesized that the mismatched stimulus would elicit Pβ1 activity. In line with our hypothesis and prior literature, we found that mismatches increase Pβ1 power but decrease ITPC. The finding of increased Pβ1 power extends prior studies showing that Pβ1 supports the encoding and maintenance of mismatched sensory information^5,11,13^. Although we did not design our task to explicitly test addeley’s WM model, our results are consistent with evidence suggesting that Pβ1 is an electrophysiological substrate of the episodic buffer^11,12^. Here, the episodic buffer may provide the space needed compare the incoming stream of information in the test sequence to the sample sequence to determine whether there was a mismatch.

Additionally, desynchronized Pβ1 activity may modulate storage capacity for sensory information, so with increased Pβ1 activity there would be a decrease in encoded information^28^. The decrease we observed in Pβ1 TPC may have served to prevent the maintenance of the mismatched stimulus. Both Pβ1 responses persisted from presentation of the test sequence into the post-test delay, consistent with the persistent activity model of WM^19–24^. Persistent activity is thought to underlie, in part, a discrete attractor network model that predicts a single outcome when there is a clear number of competing states^22,25^. In this case, there would be two competing states: match or mismatch. Our results suggest that persistent increased Pβ1 power and decreased Pβ1 TPC activity act as a discrete attractor that allows one to notice the mismatched stimulus but not maintain it. Collectively, our findings establish Pβ1 in mismatch detection, support Pβ1 as an electrophysiological substrate of the episodic buffer in WM and build support for the persistent activity framework broadly.

Finally, we explored whether mismatch responses might also be represented in a latent state during the delay, where the response is present but not visible in the EEG. We observed evidence of latent activity on mismatch trials near the end of the post-test delay. This finding is, in part, consistent with the latent activity model of WM, in which latent neural activity may help maintain unattended information that is nonetheless needed to complete the task^14,15,16,17,18^.

However, we also acknowledge that the latent effects we observed may instead be due to participants no longer needing to actively attend to the series of shapes. Rather, during the post-test delay, they only need to maintain the decisional state of match or mismatch; as such, pinging the brain may have revealed a latent decisional state. Overall, in addition to building support for the persistent activity framework, our findings reveal latent WM activity related to mismatch detection.

In summary, our EEG findings extend the role of FMθ and Pβ1 to mismatch detection. We established that FMθ activity, an electrophysiological substrate of the need for executive control and proposed substrate of the central executive, underlies successful mismatch detection and predicts behavior. However, we observed PSW ERPs to mismatches, rather than the N200, challenging a domain-general role of the N200 across the related domains of novelty, error, conflict, and mismatch processing. We further demonstrated that Pβ1 activity, which is implicated in the maintenance of sensory information and a proposed electrophysiological substrate of the episodic buffer, dissociates matched from mismatched sensory inputs. Lastly, our exploratory analyses revealed evidence of both persistent Pβ1 activity, pointing to its involvement in mismatch encoding but not maintenance, and latent activity. These findings contribute to a growing literature on error and mismatch detection ^1,2,6,7,13^ and, broadly, on how our brains transform sensory information into meaningful behavioral outputs in the service of goals^29^.

## Methods

### Participants

Thirty-five healthy adults were included in this study (20 females and 15 males; M ± SD, age: 25.26 ± 7.02 years; education: 15.15 ± 2.28 years). Post hoc sensitivity analyses indicated that this sample size provides 80% power to detect small-medium effects at two-tailed alpha < 0.05 (dependent-samples t-test Cohen’s d = 0.49; correlation n = 0.44)^30^. Participants were included based on above-chance task performance (inclusion criterion: <0.35 error in each condition, chance 0.5)^32^. Four additional participants were excluded based on this performance criterion. All participants had normal/corrected-to-normal vision and hearing, and no neurological or psychiatric diagnoses. Informed consent was obtained from all subjects. All methods were carried out in accordance with relevant guidelines and regulations. This study was approved by the University of California, Berkeley Institutional Review Board.

### Working memory delayed match-to-sample task

Participants performed our WM-DMS task, in which they detected mismatches in sequences of three colored shapes (see **Fig. 1**). On each trial, they maintained central fixation for 1.5-2.0 s (i.e., jittered intertrial interval/ITI). Then, they viewed a sample sequence of three different shapes in specific spatiotemporal positions (0.25 s stimulus, 0.25 s interstimulus interval/ISI). Each shape appeared in one of four different spatial positions (left, right, top, bottom), followed by a 2-s delay and one of four conditions of test sequence with identical timing to the sample sequence. The test sequence matched the sample sequence exactly (i.e., match) or was changed on one dimension: two out of three stimuli were either new (mismatch: identity), swapped across spatial positions (mismatch: spatial), or swapped across temporal positions (mismatch: temporal). On mismatch trials, one stimulus always matched on all three dimensions, and the temporal positions of the two mismatched stimuli were evenly counterbalanced (i.e., 1 and 2, 2 and 3, 1 and 3; see **Fig. 1A**). Following a second 2-s delay, they responded match/mismatch/unsure by self-paced mouse click. The central fixation crosshair remained on screen for the duration of the task. Following eight practice trials, participants completed 256 trials with a break every 16 trials. The four test conditions were evenly counterbalanced and presented in pseudo-randomized order with four trials per condition every 16 trials. For half of all trials, a star flashed for 0.1 s midway through both delays in the center of the screen at randomly jittered times ranging from 1.0-1.15 s from the onset of the delay (see **Fig. 1C**). The star trials were evenly counterbalanced across conditions and appeared on eight of every 16 trials. The task was programmed in MATLAB (MathWorks Inc., Natick, MA) with the Psychtoolbox-3 software extension.

Behavioral accuracy rate was calculated per participant as the hit rate (i.e., proportion of match sequences that were correctly identified as match) minus false alarm rate (proportion of mismatched sequences that were incorrectly identified as match). This procedure defines chance accuracy as zero and corrects for differences in trial counts between conditions as well as an individual’s tendency to respond match/mismatch^33^. A 4 (condition: match, mismatch identity, mismatch spatial, mismatch temporal) × 2 (star, no star) within-subjects ANOVA on accuracy (proportion correct) indicated that neither the star/no star main effect nor interaction with condition was significant (p > 0.45). This analysis also revealed an expected main effect of condition (mismatch > match; F(3,279) = 27.46, p < 0.001), which qualified our decision to analyze accuracy using the corrected measure of accuracy rate. RT was calculated per participant across all trials.

### EEG data acquisition and preprocessing

EEG data were acquired at a sampling rate of 1,024 Hz using a 64 + 8 channel BioSemi ActiveTwo amplifier with Ag-AgCl pin-type active electrodes mounted on a cap based on the extended 10-20 system (BioSemi, Amsterdam, NL). The horizontal electrooculogram (EOG) was recorded at both external canthi and the vertical EOG was monitored with a right inferior eye electrode. Two electrodes were placed on the earlobes for offline referencing. Electrode impedances were kept below 20 kΩ. Raw data were bandpass (0.1-100 Hz finite impulse response filter) and notch filtered (60-Hz line noise removed using discrete Fourier transform), demeaned, and separated into 8.5-s trials (−1 s from the onset of the sample sequence to +1 s from the offset of the post-test delay). Filtered data were down sampled to 512 Hz and any artifactual channels (e.g., from signal dropout) were removed. Independent components analysis was performed on the remaining channels to remove EOG, electromyography, and other artifacts^34^. Any channels removed were replaced with the interpolated mean values from neighboring channels (mean 7.6 channels). Data were manually inspected blind to task parameters to remove trials containing residual noise. The surface Laplacian spatial filter was applied to minimize volume conduction and enhance the source signal ^35–37^. Finally, incorrect trials were excluded^32,38,39,40,41,42,43,44^, resulting in a mean of 203.66 (SD: 27.22) artifact-free, correct trials analyzed per participant. EEG preprocessing and analysis routines utilized functions from the open-source FieldTrip toolbox for MATLAB^45^..

### EEG data quantification

#### Event-related potentials

The 8.5-s data segments were zero-padded to 15 s to minimize filtering-induced edge artifacts, lowpass filtered at 20 Hz (finite impulse response filter)^46^, detrended, and separated into epochs corresponding to the pre-stimulus baseline (−0.75 to -0.25 s from sample onset) and test sequence (3.25-4.5 s from sample onset). Test sequences were realigned per trial so that the zero point indicated the onset of the first mismatched stimulus (i.e., first or second shape in the sequence), and then cut into 1-s epochs (0-1s from stimulus onset; see **Fig. 1A**). Matched test sequences were submitted to the same realignment procedure, with the zero point randomly assigned to the second stimulus on one third of trials to account for any position effects. ERPs were absolute baseline-corrected (i.e., realigned test sequence – baseline mean) and averaged across each of the four test conditions. ERP data were then averaged across the three mismatch conditions for comparison against the match condition.

#### Spectral decomposition

The 8.5-s data segments were zero-padded to 15 s to minimize filtering-induced edge artifacts and the Hanning taper time-frequency spectrum was calculated by sliding a frequency-dependent window of three cycles (Δt = /f) in 0-ms steps at each frequency from 2.5-50 Hz (0.5-Hz resolution, 2-Hz bandwidth). Test sequences were realigned using the same procedure described above (see *Event-Related Potentials*). Power and phase clustering (i.e., intertrial phase coherence/ITPC) were computed from the amplitude and phase outputs of the time-frequency spectrum, respectively. Power was relative-change baseline-corrected (i.e., (realigned test sequence – baseline mean)/baseline mean) and averaged across each of the four test conditions. ITPC was computed for each of the four test conditions. Power and ITPC data were then averaged across the three mismatch conditions for comparison against the match condition.

#### Exploratory analysis of the post-test delay

ERP, power, and ITPC data were recomputed and averaged separately across conditions for star- and no-star trials, following the same procedures described above. To explore persistent representations, the no-star trial data were cut into 2.5-s epochs (−0.5-2 s from delay onset, including the second ITI and third stimulus in the test sequence; see **Fig. 1B**), with no realignment. To explore latent representations, the star trial data were realigned per trial such that the zero point indicated the onset of the star and then cut into 1.85-s epochs (−1 to 0.85 s from star onset; see **Fig. 1C**).

### Statistics

EEG signatures of mismatch detection were identified using dependent-samples Monte Carlo permutations on all 64 channels, with cluster-based correction for multiple comparisons^47^.

Clusters were formed in space, time, and frequency (for power and ITPC data) by thresholding t-values at p < 0.05 using the maximum sum criterion. Permutation distributions were generated by randomly shuffling match/mismatch data labels (1,000 iterations) and corrected p-values were obtained by comparing the observed data to the random permutation distributions. This is an extremely powerful approach because it recreates any biases in the data with each randomization and tests for mismatch effects without any assumptions about their spatial, temporal, or spectral distribution. We also analyzed differences between all four conditions using an F-test on all 64 channels, with the same cluster-based correction for multiple comparisons. Cluster-based statistics were conducted using the FieldTrip toolbox for MATLAB^45^.

Data-driven post hoc correlations examined relationships between EEG mismatch detection effects (e.g., frontal midline ERP and FMθ effects), and between EEG mismatch detection effects and individual behavioral outcomes (i.e., accuracy rate, RT). Spearman’s rank correlation coefficients were calculated from mismatch versus match differences averaged across cluster-corrected data points^39^.

## Acknowledgements and Funding

We thank Vasanth Kommu for helping collect data, and members of the Northwestern University Dynamic Brain Lab for their crucial feedback. The work in this paper was funded by NIH grants R00NS115918, R01NS021135, and T32NS047987. This research was supported in part through the computational resources and staff contributions provided for the Quest high performance computing facility at Northwestern University which is jointly supported by the Office of the Provost, the Office for Research, and Northwestern University Information Technology. Funding sources were not involved in the study design; data collection, analysis, and interpretation; writing of the report; or publication submission.

## Author Contributions

**J.Y**.: Conceptualization, Validation, Formal analysis, Data Curation, Writing – Original Draft, Writing - Review & Editing, Visualization **L.S**.: Formal analysis, Data Curation, Writing - Review & Editing **K.C**.: Investigation, Writing - Review & Editing **R.K**.: Conceptualization, Resources, Writing - Review & Editing, Project Administration, Funding acquisition **E.J**.: Conceptualization, Methodology, Software, Formal analysis, Investigation, Resources, Data Curation, Writing - Review & Editing, Visualization, Supervision, Project Administration, Funding acquisition

## Data Availability

The datasets generated and/or analyzed during the current study are available in the OSF repository, https://osf.io/y7dj6/.

## Ethics

This study was approved by the University of California, Berkeley Institutional Review Board.

## References

1. Cavanagh, J. F. & Frank, M. J. Frontal theta as a mechanism for cognitive control. Trends in Cognitive Sciences vol. 18 414–421 Preprint at 10.1016/j.tics.2014.04.012 (2014).

2. Polich, J. Updating P300: An integrative theory of P3a and P3b. Clinical Neurophysiology vol. 118 2128–2148 Preprint at 10.1016/j.clinph.2007.04.019 (2007).

3. Gelastopoulos, A., Whittington, M. A. & Kopell, N. J. Parietal low beta rhythm provides a dynamical substrate for a working memory buffer. Proc Natl Acad Sci U S A 116, 16613–16620 (2019).

4. Michels, L. et al. Simultaneous EEG-fMRI during a working memory task: Modulations in low and high frequency bands. PLoS One 5, (2010).

5. Roberts, B. M., Hsieh, L. T. & Ranganath, C. Oscillatory activity during maintenance of spatial and temporal information in working memory. Neuropsychologia 51, 349–357 (2013).

6. Cavanagh, J. F., Zambrano-Vazquez, L. & Allen, J. J. B. Theta lingua franca: A common mid-frontal substrate for action monitoring processes. Psychophysiology 49, 220–238 (2012).

7. Folstein, J. R. & Van Petten, C. Influence of cognitive control and mismatch on the N2 component of the ERP: A review. Psychophysiology 152–170 (2008) doi:10.1111/j.1469-8986.2007.00602.x.

8. Itthipuripat, S., Wessel, J. R. & Aron, A. R. Frontal theta is a signature of successful working memory manipulation. Exp Brain Res 224, 255–262 (2013).

9. Maurer, U. et al. Frontal Midline Theta Reflects Individual Task Performance in a Working Memory Task. Brain Topogr 28, 127–134 (2015).

10. Baddeley, A. D. & Hitch, G. Working Memory. (1974) doi:10.1016/S0079-7421(08)60452-1.

11. Baddeley, A. The episodic buffer: a new component of working memory? Trends Cogn Sci 4, 417–423 (2000).

12. Baddeley, A. D., Allen, R. J. & Hitch, G. J. Binding in visual working memory: The role of the episodic buffer. Neuropsychologia 49, 1393–1400 (2011).

13. Haenschel, C., Baldeweg, T., Croft, R. J., Whittington, M. & Gruzelier, J. Gamma and beta frequency oscillations in response to novel auditory stimuli: A comparison of human electroencephalogram (EEG) data with in vitro models. Proceedings of the National Academy of Sciences 97, 7645–7650 (2000).

14. Barbosa, J. et al. Interplay between persistent activity and activity-silent dynamics in the prefrontal cortex underlies serial biases in working memory. Nat Neurosci 23, 1016–1024 (2020).

15. Hedayati, S., O’Donnell, R. E. & Wyble,. model of working memory for latent representations. Nat Hum Behav 6, 709–719 (2022).

16. Sprague, T. C., Ester, E. F. & Serences, J. T. Restoring Latent Visual Working Memory Representations in Human Cortex. Neuron 91, 694–707 (2016).

17. Stokes, M. G. ‘ctivity-silent’ working memory in prefrontal cortex: dynamic coding framework. Trends in Cognitive Sciences vol. 19 394–405 Preprint at 10.1016/j.tics.2015.05.004 (2015).

18. Wolff, M. J., Ding, J., Myers, N. E. & Stokes, M. G. Revealing hidden states in visual working memory using electroencephalography. Front Syst Neurosci 9, (2015).

19. Curtis, C. E. & Sprague, T. C. Persistent Activity During Working Memory From Front to Back. Frontiers in Neural Circuits vol. 15 Preprint at 10.3389/fncir.2021.696060 (2021).

20. Funahashi, S., Bruce, C. J. & Goldman-Rakic, P. S. Mnemonic Coding of Visual Space in e nkey’ D l e l P ef n l C ex. JOURNALOFNEUROPHYSIOLOGY vol. 6 (1989).

21. Goldman-Rakic, P. S. Cellular Basis of Working Memory Review. Neuron vol. 14 (1995).

22. Inagaki, H. K., Fontolan, L., Romani, S. & Svoboda, K. Discrete attractor dynamics underlies persistent activity in the frontal cortex. Nature 566, 212–217 (2019).

23. Li, N., Daie, K., Svoboda, K. & Druckmann, S. Robust neuronal dynamics in premotor cortex during motor planning. Nature 532, 459–464 (2016).

24. Romo, R., Brody, C., Hernandez, A. & Lemus, L. Neuronal correlates of parametric working memory in the prefrontal cortex. Nature 470–473 (1999) doi:10.1038/20939.

25. Kamiński, J. & Rutishauser, U. etween persistently active and activity-silent frameworks: novel vistas on the cellular basis of working memory. Annals of the New York Academy of Sciences vol. 1464 64–75 Preprint at 10.1111/nyas.14213 (2020).

26. De Gee, J. W., Correa, C. M. C., Weaver, M., Donner, T. H. & Van Gaal, S. Pupil Dilation and the Slow Wave ERP Reflect Surprise about Choice Outcome Resulting from Intrinsic Variability in Decision Confidence. Cerebral Cortex 31, 3565–3578 (2021).

27. García-Larrea, L. & Cézanne-Bert, G. P3, Positive slow wave and working memory load: a study on the functional correlates of slow wave activity. Electroencephalogr Clin Neurophysiol 108, 260–273 (1998).

28. Miller, E. K., Lundqvist, M. & Bastos, A. M. Working Memory 2.0. Neuron vol. 100 463–475 Preprint at 10.1016/j.neuron.2018.09.023 (2018).

29. Christophel, T. B., Klink, P. C., Spitzer, B., Roelfsema, P. R. & Haynes, J. D. The Distributed Nature of Working Memory. Trends in Cognitive Sciences vol. 21 111–124 Preprint at 10.1016/j.tics.2016.12.007 (2017).

30. Faul, F., Erdfelder, E.Lang, A.-G. & Buchner, A. G*Power 3: A flexible statistical power analysis program for the social, behavioral, and biomedical sciences. Behav Res Methods 39, 175–191 (2007).

31. Faul, F., Erdfelder, E., Buchner, A. & Lang, A.-G. Statistical power analyses using G*Power 3.1: Tests for correlation and regression analyses. Behav Res Methods 41, 1149–1160 (2009).

32. Johnson, E. L. et al. A rapid theta network mechanism for flexible information encoding. Nat Commun 14, (2023).

33. Snodgrass, J. G. & Corwin, J. Pragmatics of Measuring Recognition Memory: Applications to Dementia and Amnesia. Journal of Experimental Psychology: General vol. 117 (1988).

34. Hipp, J. F. & Siegel, M. Dissociating neuronal gamma-band activity from cranial and ocular muscle activity in EEG. Front Hum Neurosci (2013) doi:10.3389/fnhum.2013.00338.

35. Cohen, M. X. Comparison of different spatial transformations applied to EEG data: A case study of error processing. International Journal of Psychophysiology 97, 245–257 (2015).

36. He, B., Sohrabpour, A., Brown, E. & Liu, Z. Electrophysiological Source Imaging: A Noninvasive Window to Brain Dynamics. Annu Rev Biomed Eng 171–196 (2018) doi:10.1146/annurev-bioeng-062117.

37. Lai, M., Demuru, M., Hillebrand, A. & Fraschini, M. A comparison between scalp- and source-reconstructed EEG networks. Sci Rep 8, (2018).

38. Jones, K. T., Peterson, D. J., Blacker, K. J. & Berryhill, M. E. Frontoparietal neurostimulation modulates working memory training benefits and oscillatory synchronization. Brain Res 1667, 28–40 (2017).

39. Johnson, E. L. et al. Orbitofrontal cortex governs working memory for temporal order. Current Biology 32, R410–R411 (2022).

40. Johnson, E. L. et al. Spectral imprints of working memory for everyday associations in the frontoparietal network. Front Syst Neurosci 12, (2019).

41. Johnson, E. L. et al. Dynamic frontotemporal systems process space and time in working memory. PLoS Biol 16, (2018).

42. Johnson, E. L. et al. Bidirectional Frontoparietal Oscillatory Systems Support Working Memory. Current Biology 27, 1829-1835.e4 (2017).

43. Davoudi, S., Dezfouli, M. P., Knight, R. T., Daliri, M. R. & Johnson, E. L. Prefrontal lesions disrupt posterior alpha–gamma coordination of visual working memory representations. J Cogn Neurosci 33, 1798–1810 (2021).

44. Parto Dezfouli, M., Davoudi, S., Knight, R. T., Daliri, M. R. & Johnson, E. L. Prefrontal lesions disrupt oscillatory signatures of spatiotemporal integration in working memory. Cortex 138, 113–126 (2021).

45. Oostenveld, R., Fries, P., Maris, E. & Schoffelen, J. M. FieldTrip: Open source software for advanced analysis of MEG, EEG, and invasive electrophysiological data. Comput Intell Neurosci 2011, (2011).

46. Kappenman, E. S., Farrens, J. L., Zhang, W., Stewart, A. X. & Luck, S. J. ERP CORE: An open resource for human event-related potential research. Neuroimage 225, 117465 (2021).

47. Maris, E. & Oostenveld, R. Nonparametric statistical testing of EEG- and MEG-data. J Neurosci Methods 164, 177–190 (2007).

